# Plaques formed by mutagenized viral populations have elevated co-infection frequencies

**DOI:** 10.1101/108092

**Authors:** Elizabeth R. Aguilera, Andrea K. Erickson, Palmy R. Jesudhasan, Christopher M. Robinson, Julie K. Pfeiffer

**Author notes:** Corresponding Author: Julie K. Pfeiffer.

## Abstract

The plaque assay is a common technique used to measure virus concentrations and is based upon the principle that each plaque represents a single infectious unit. As such, plaque number is expected to correlate linearly with the virus dilution plated and each plaque should be formed by a single founder virus. Here, we examined whether more than one virus can contribute to plaque formation. By using genetic and phenotypic assays with genetically marked polioviruses, we found that multiple parental viruses are present in 5-7% of plaques, even at an extremely low multiplicity of infection. We demonstrated through visual and biophysical assays that, like many viral stocks, our viral stocks contain both single particles and aggregates. These data suggest that aggregated virions are capable of inducing co-infection and chimeric plaque formation. In fact, inducing virion aggregation via exposure to low pH increased co-infection in a flow cytometry-based assay. We hypothesized that plaques generated by viruses with high mutation loads may have higher co-infection frequencies due to fitness restoring processes such as complementation and recombination. Indeed, we found that co-infection frequency correlated with mutation load, with 17% chimeric plaque formation for heavily mutagenized viruses. Importantly, the frequency of chimeric plaques may be underestimated by up to three-fold, since co-infection with the same parental virus cannot be scored in our assay. This work indicates that more than one virus can contribute to plaque formation and that co-infection may assist plaque formation in situations where the amount of genome damage is high.

## IMPORTANCE

One of the most common methods to quantify viruses is the plaque assay, where it is generally presumed that each plaque represents a single infectious virus. Using genetically marked polioviruses, we demonstrate that a plaque can contain more than one parental virus, likely due to aggregates within virus stocks that induce co-infection of a cell. A relatively small number of plaques are the products of co-infection for our standard virus stocks. However, mutagenized virus stocks with increased genome damage give rise to more “chimeric” plaques. These results suggest that co-infection may aid plaque formation of viruses with genome damage, possibly due to processes such as complementation and recombination. Overall, our results suggest that the relationship between viral dilution and plaque number may not be linear, particularly for mutagenized viral populations.

## INTRODUCTION

Viral concentrations are frequently determined using the plaque assay, which is based on the principle that each plaque represents one infectious unit (1). Many mammalian viruses have high particle to plaque-forming unit (PFU) ratios due to various factors such as assembly defects, mutations, and inefficient steps during the viral replication cycle (2–4). For most mammalian viruses, the number of plaques is directly proportional to the concentration of virus. This indicates that one infectious particle gives rise to one plaque, providing a “one hit” model for plaque formation. However, certain viruses of plants and fungi have a “two hit” model, whereby two particles containing different genome segments are required to co-infect the same cell to facilitate productive infection and plaque formation (5,6). Recently, Ladner *et al*. reported that a mammalian virus has a “three hit” model (7). This virus is composed of five genome segments, each packaged in five separate viral particles; however, a minimum of three segments were required to facilitate productive infection. While previous work indicates that, for most mammalian viruses, plaque number appears to correlate linearly with the dilution of virus plated, it is possible that some plaques may be the products of co-infection.

Several mechanisms could facilitate co-infection of viruses. Previous reports demonstrated that cells may be infected by more than one virus at a higher frequency than predicted by Poisson distribution (8–12). For example, poliovirus can be packaged in phosphatidylserine vesicles, which promotes co-infection of neighboring cells (8). Multiple coxsackievirus B3 and hepatitis A virions can also be packaged in vesicle-like structures (13,14). Vesicular stomatitis virus was found to form plaques containing two different viral genomes, indicating that co-infection occurred (9). Additionally, many different viruses aggregate in solution and could induce co-infection (15–18). Viral stocks of vaccinia virus, influenza virus, adenovirus, herpes virus, and echovirus contained virion aggregates that were resistant to antibody-mediated neutralization and/or radiation (18,19). Poliovirus and reovirus particles can aggregate in sewage, which may contribute to initial infection of the host and it is possible that virion aggregates exist *in vivo* (20,21).

RNA viruses undergo error-prone replication due to lack of proofreading activity of their RNA-dependent RNA polymerase (RdRp). For poliovirus, the mutation frequency is ~10^-4^ per nucleotide per replication cycle (22). Mutations can be beneficial to viruses in some circumstances, for example by conferring resistance to neutralizing antibodies. However, most mutations are deleterious and reduce viral fitness (23). Much has been learned about RNA virus mutation-associated fitness effects from viral populations harboring increased or decreased mutation frequencies. Crotty *et al*. demonstrated that the nucleoside analog ribavirin (RBV) is an RNA virus mutagen. RBV is incorporated into nascent RNA by the viral RdRp, which increases transition mutations and can cause “error catastrophe” (24). Conversely, poliovirus passaged in the presence of RBV acquired a single point mutation in the RdRp, G64S, which increased fidelity of viral RNA synthesis and reduced the error frequency of the viral population (25,26). Importantly, G64S poliovirus had reduced viral fitness during infection of mice, indicating that viral population diversity is necessary for virulence (27,28).

RNA viruses may overcome mutation-induced fitness costs by several genetic mechanisms (23). First, deleterious mutations may revert via error prone RNA synthesis. Second, genetic recombination can generate progeny genomes lacking deleterious mutations. Recombination can occur when two viruses co-infect the same cell and exchange genetic information, likely through a copy-choice mechanism (29). Recombination of genomes has been observed in poliovirus and other enteroviruses (30–36). Furthermore, defective RNA genomes are capable of undergoing recombination *in vivo*, thus restoring their fitness (36). Third, fitness may be restored by complementation, whereby viruses with distinct genetic defects complement one another. Complementation has been observed within brain tissues of poliovirus-infected mice (28). Fourth, reassortment can occur when two distinct segmented viruses co-infect the same cell and generate progeny with genome segments from both viruses. Unlike reversion, fitness restoration by recombination, complementation, and reassortment generally requires synchronous co-infection of a cell with more than one virus. Overall, these genetic processes can alter viral diversity and increase fitness.

In this work, we examined whether viral plaques are derived from a single founder and whether viruses with increased genome damage may be more reliant on co-infection for plaque formation. Through the use of a genetic assay with ten distinct polioviruses and a phenotypic assay with two distinct polioviruses, we have shown that multiple parental viruses can be found within a single plaque, which we refer to as a chimeric plaque. We determined that 5-7% of plaques were derived from two or more viruses. To assess factors contributing to co-infection, we used dynamic light scattering and electron microscopy to demonstrate that viral stocks contain both single particles and aggregates, suggesting that infection with virion aggregates is likely responsible for chimeric plaque formation. Indeed, inducing virion aggregation via exposure to low pH increased co-infection frequency in a separate flow cytometry-based assay. We examined whether there were situations where co-infection frequencies varied and whether co-infection may assist plaque formation. Using high fidelity G64S polioviruses that harbor fewer mutations than wild-type (WT) and RBV-mutagenized polioviruses that harbor more mutations than WT, we found that co-infection frequency correlated with mutation load. In fact, 17% of plaques from mutagenized virus infections were the product of co-infection. This work indicates more than one virus can contribute to plaque formation and that co-infection may assist plaque formation in situations where the amount of genome damage is high.

## RESULTS

### A small percentage of plaques are derived from more than one parental virus

To determine whether plaques are the product of more than one founding virus, distinct parental viruses that can be discriminated by genotype or phenotype are required. We began by using ten genetically distinct polioviruses that each contain unique silent point mutations and are discriminated by hybridization of RT-PCR products with specific probes. We previously demonstrated that these ten marked viruses are equally fit (37–39). We mixed equal amounts of the ten viruses and infected HeLa cells at an extremely low MOI such that ~2-20 plaques would be generated on each plate of 10^6^ cells (Fig. 1A). Plaques were picked, viruses were amplified for a single cycle in HeLa cells, and the presence of each virus was determined by probe-specific hybridization of RT-PCR products (37–39). Plaques were scored as having a single parent virus (e.g. Plaque 1) or more than one parent virus (e.g. Plaque 2) (Fig. 1B). We examined whether each of the ten viruses were equally represented in plaques since skewed ratios of the input viruses could impact the observed frequency of co-infection. The distribution of viruses present within all plaques tested was reasonably even, with each of the ten viruses nearly equally represented (Fig. 1C). Of 123 total plaques analyzed, 6 had more than one founding parental viruses and 117 had a single virus (Table 1). Therefore, 4.9% of plaques were derived from more than one parental virus.

**Table 1.** **Frequency of plaques with more than one founding/parental virus.**

| **<u>System</u>** | **<u>Fraction with > 1 virus</u>** | **<u>Percent with > 1 virus</u>** |
| --- | --- | --- |
| 10 Virus<br>Genotypic Assay<sup>a</sup> | 6/123 | 4.9% |
| 2 Virus<br>Phenotypic Assay<sup>b</sup> | 10/138 | 7.3% |

<sup>a</sup> Assay described in Figure 1
<sup>b</sup> Assay described in Figure 2

**Figure 1.**
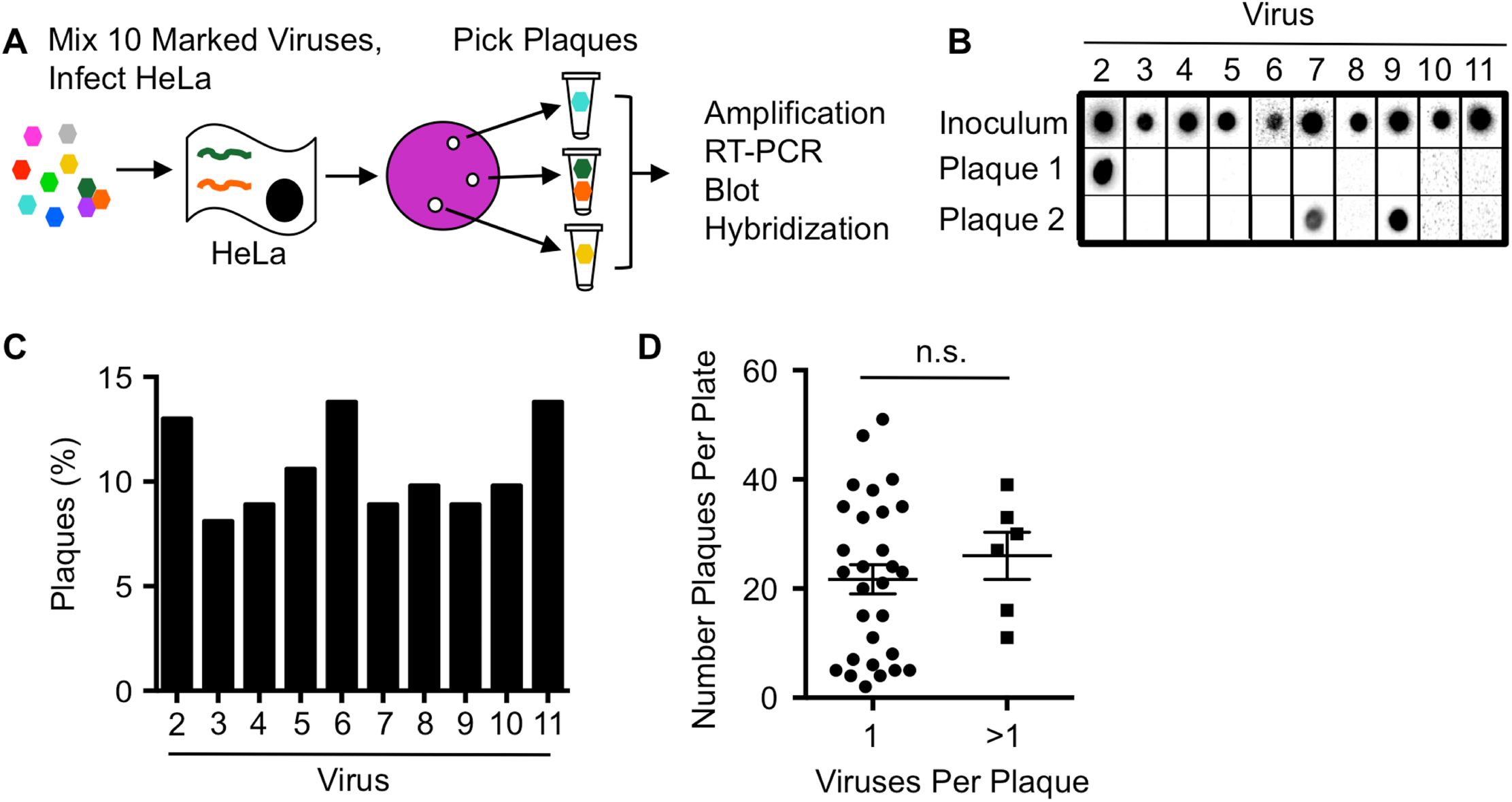
Genotypic assay reveals co-infection of polioviruses. (A) Schematic of assay design. HeLa cells were infected at an MOI of ~0.00001 with an equal mixture of ten genetically marked viruses. Hypothetical genomes are depicted in a HeLa cell. Plaques were picked from the agar overlays after incubation at 37°C for 48 h. Plaque viruses were amplified by infecting new cells and RT-PCR products were blotted and probed on a membrane to identify the virus(es) present. (B) Representative plaque virus samples detected by probes. The number of viruses present within each plaque was quantified. (C) Distribution of the ten marked viruses among all plaques. (D) Frequency of co-infected or non co-infected plaque viruses versus the number of plaques per plate, with mean ± standard error of the mean (difference not significant [n.s.], determined by Student’s *t*-test).

It was possible that overlapping plaques with single parental viruses were picked and incorrectly scored as chimeric. If so, the frequency of chimeric plaques should be higher on plates with more plaques present. To rule out the possibility that picking dual-parent plaques was enriched in situations where a higher number of plaques were present on the plate, we compared the number of plaques on plates from single and dual parent plaques (Fig. 1D). We found that dual parent plaques were not more prevalent on plates with higher plaque numbers, indicating that cross-contamination of plaque viruses was unlikely.

Since the ten virus genotypic assay to measure co-infection is relatively labor intensive, we sought to simplify the screening process by using a previously characterized poliovirus mutant that can be discriminated from WT by a phenotypic assay (29) (Fig. 2A). *3NC-202gua*^*R*^ poliovirus has mutations that confer resistance to guanidine hydrochloride and temperature sensitivity (called Drug^R^/Temp^S^ hereafter). Guanidine is used as a protein denaturant but can also specifically inhibit poliovirus RNA synthesis (40). Conversely, WT poliovirus is guanidine sensitive and temperature resistant (called Drug^S^/Temp^R^ hereafter). We mixed equal amounts of Drug^R^/Temp^S^ and Drug^S^/Temp^R^ polioviruses and infected HeLa cells at an extremely low MOI such that ~2-20 plaques would be generated on each plate of 10^6^ cells. Infections were incubated in permissive conditions for both parental viruses (33°C without drug). Plaques were picked and analyzed by their growth phenotypes to determine whether one or both parental viruses were present. Plaque formation at 33°C in the presence of drug indicated the presence of the Drug^R^/Temp^S^ parent. Plaque formation at 39°C indicated the presence of the Drug^S^/Temp^R^ parent. Plaque formation in both conditions indicated presence of both parental viruses (Fig. 2C). The distribution of viruses present within plaques was reasonably even, with 44.2% and 55.8% of Drug^S^/Temp^R^ and Drug^R^/Temp^S^ parental viruses, respectively (Fig. 2C). In this assay, we found that 10 out of 138 (7.3%) plaques had more than one parental virus present (Table 1). Therefore, both the genotypic and phenotypic assays indicate that ~5-7% of plaques that arise following infection of HeLa cells contain more than one distinct parental virus.

**Figure 2.**
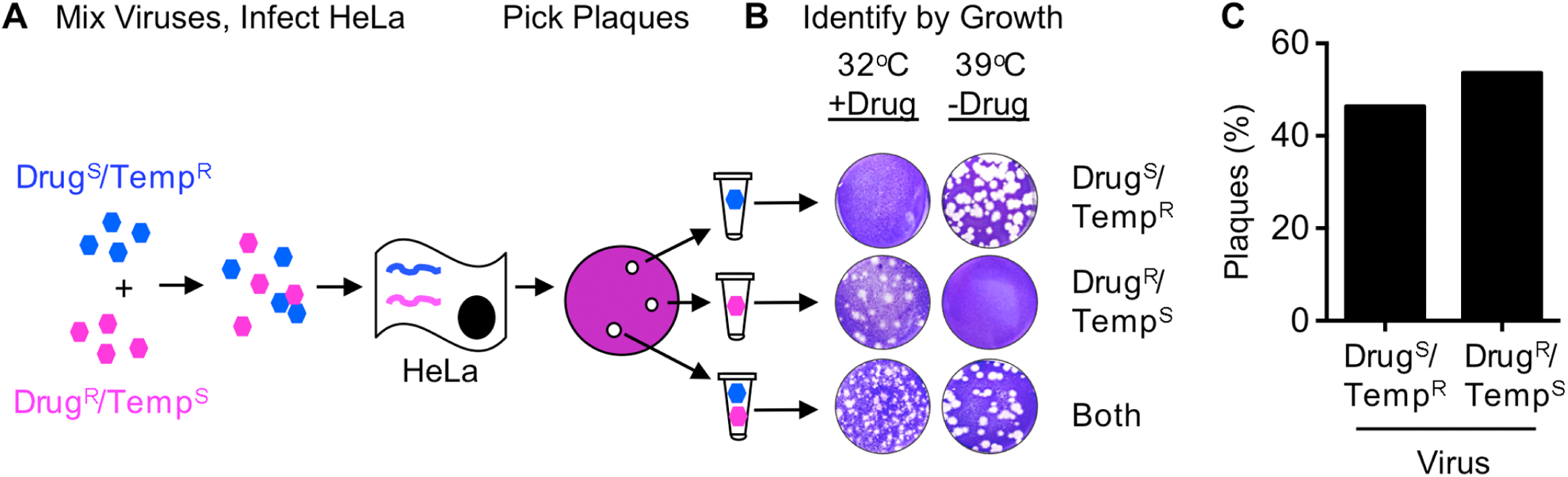
Phenotypic assay reveals co-infection of polioviruses. (A) Schematic of co-infection assay using Drug^S^/Temp^R^ and Drug^R^/Temp^S^ viruses. The two parental viruses were mixed, incubated and HeLa cells were infected with the viral mixture at an MOI of ~0.00001. Hypothetical viral genomes are depicted in a HeLa cell. Plaques were picked 4-5 days after adding agar overlay at 33°C in the absence of guanidine (permissive conditions). (B) Representative plaques in the phenotypic scoring assay. Plaque viruses were plated on HeLa cells under dual selective conditions as indicated. (C) Distribution of the two parental viruses among all plaques.

### Poliovirus stocks contain viral aggregates

To examine whether co-infection of poliovirus may be due to viral aggregation, we examined representative viral stocks using visual and biophysical assays. We first examined viruses using electron microscopy and observed single viral particles as well as aggregated viral particles (Fig. 3A), in agreement with previous studies (15–17). To quantify poliovirus aggregation, we used dynamic light scattering, which measures the size of particles in solution. Using this assay, we observed that our viral stock contained both single particles (15 nm radius) and aggregates ranging from 2-10 particles (Fig. 3B). Overall, these results indicate that viral stocks contain aggregates and we speculated that virion aggregation facilitates co-infection of viruses.

**Figure 3.**
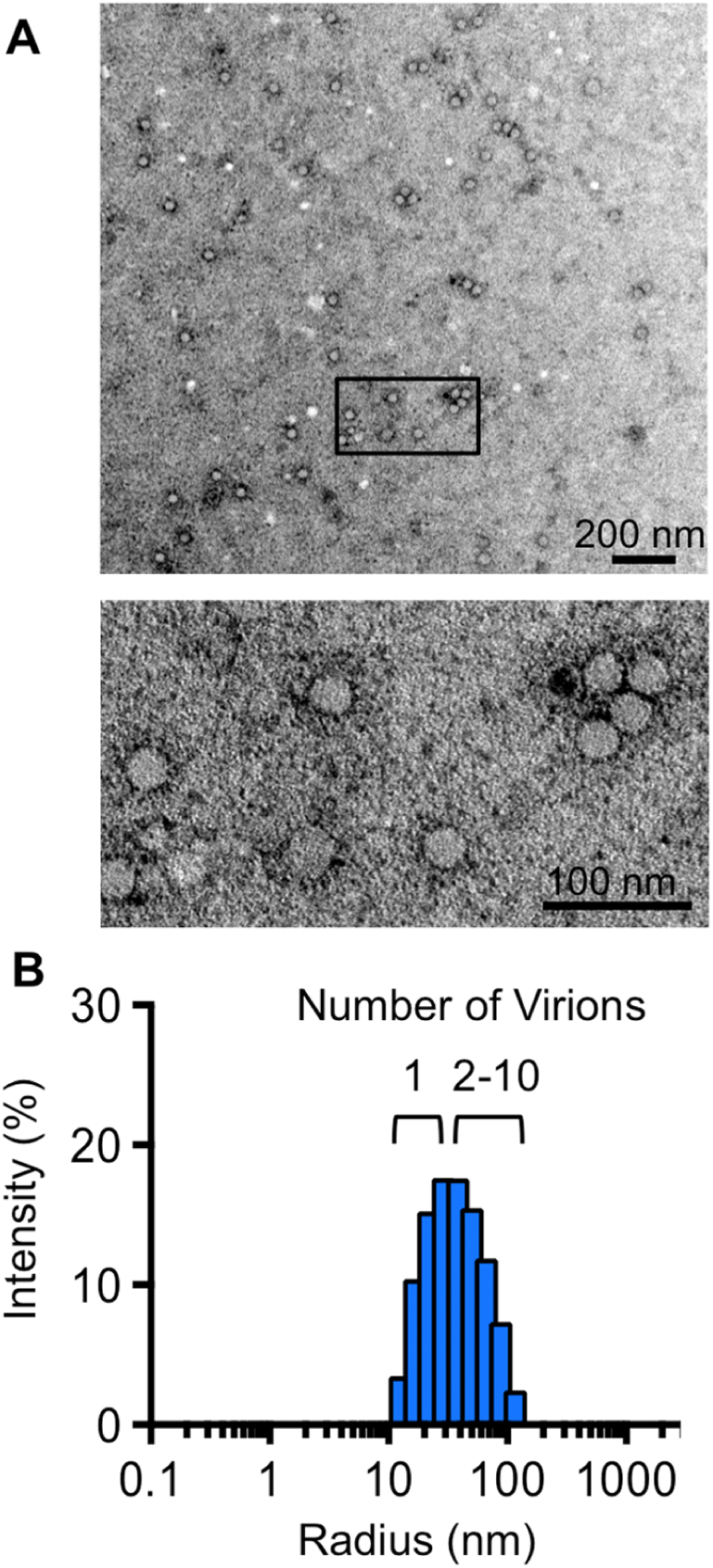
Stocks of poliovirus contain aggregates. (A) Transmission electron microscopy of a representative poliovirus stock. Viral particles were imaged at a magnification of 13,000x (top image) or 30,000x (bottom image, a detail of the boxed region in the top image). (B) Dynamic light scattering analysis of a representative poliovirus stock. Virus stock was diluted to 5 × 10^4^ PFU/mL and centrifuged for 10 min prior to analysis on a Protein Solutions DynaPro instrument. Poliovirus radius = 15 nm.

### Inducing aggregation increases co-infection frequency

To test the hypothesis that virion aggregation facilitates co-infection, we examined co-infection frequency for virions exposed to conditions that induce aggregation using a minimally labor-intensive flow cytometry-based assay. Polioviruses expressing either GFP or DsRed (41) were mixed and incubated for 4 h in PBS (control) or in glycine-HCL buffer, pH 3, which induces aggregation of poliovirus (15–17) (Fig. 4A). Viruses were then used to infect HeLa cells at an MOI of 0.01, such that ~99% of cells remain uninfected. Sixteen hours post-infection, cells were subjected to flow cytometry to quantify the percentage of uninfected, singly infected (red or green), or co-infected cells (red and green, dual positive). To ensure that the low pH treatment induced virion aggregation, we measured particle size using dynamic light scattering. Indeed, viruses exposed to low pH had increased radii compared with viruses exposed to PBS, confirming that low pH induced virion aggregation (Fig. 4B). Based upon an MOI of 0.01, approximately 1% of cells infected with PBS-treated viruses were infected with a single virus (Fig. 4C). However, 0.83% of cells infected with low pH-treated viruses were infected with a single virus, suggesting that virion aggregation slightly reduced the total number of infectious units. A small percentage, 0.0048%, of cells infected with PBS-treated viruses were co-infected with both DsRed− and GFP-expressing viruses (Fig. 4D), which is close to the predicted number of co-infected cells based on Poisson distribution (0.005%). Interestingly, co-infection was increased 2.6-fold in low pH-treated viruses, suggesting that aggregation enhances co-infection. Furthermore, these data indicate that co-infection can occur during infections performed in liquid culture.

**Figure 4.**
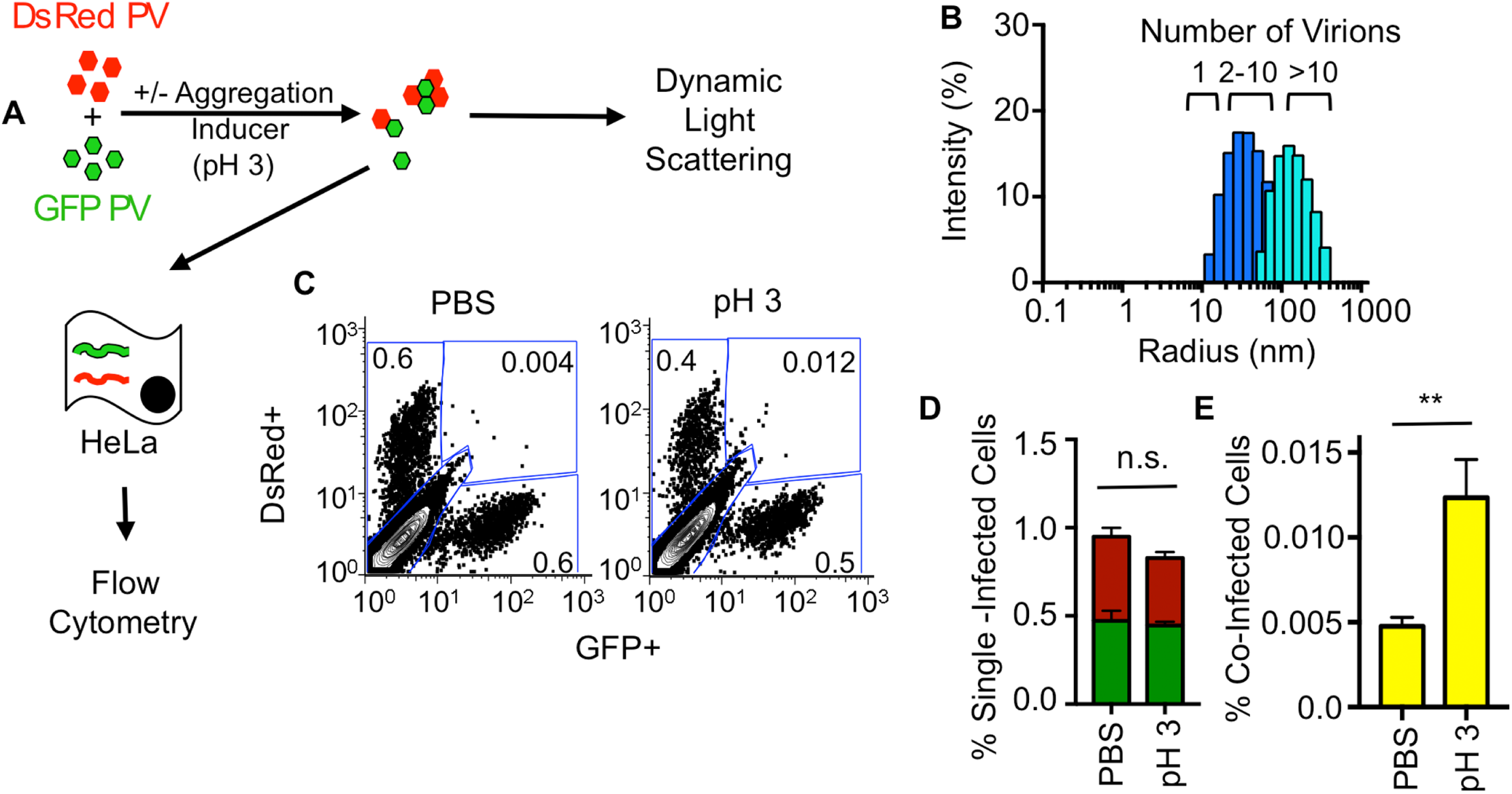
Flow cytometry-based assay demonstrates correlation between aggregation and co-infection. (A) Schematic of flow cytometry-based assay. GFP-and DsRed-expressing polioviruses were mixed in the presence or absence of aggregation inducing conditions (+/- exposure to pH 3 solution for 4 h) prior to analysis by dynamic light scattering or infection of HeLa cells at an MOI of 0.01. At 16 hpi, infection was quantified using flow cytometry. (B) Dynamic light scattering analysis of viruses exposed to PBS (royal blue, same data as in Fig. 3B) or viruses exposed to pH 3 glycine-HCL buffer for 4 h (turquoise). Samples were processed as described in Fig. (C) Representative FACS plots showing quantification of DsRed, GFP, or dual positive cells. The units for x-and y-axes are GFP and DsRed fluorescence intensity, respectively. The numbers in each gate indicate the percentage positive of the total cell population of 2 × 10^5^ cells counted. Gates were drawn from FACS plots of HeLa cells exposed to pH 3 glycine-HCL in the absence of PV (lower left gate), infected with 1 × 10^4^ PFU GFP-PV (lower right gate), infected with 1 × 10^4^ PFU DsRed-PV (upper left gate) or infected with 1 × 10^4^ PFU GFP-PV and 1 × 10^4^ PFU DsRed-PV (upper right gate) (D) Percentage of cells infected by single viruses (GFP or DsRed). (E) Percentage of co-infected cells, positive for both GFP and DsRed (upper right gate). Results are presented as mean ± standard error of the mean (n=9). Statistical significance was determined by Student’s *t*-test, \*\**P*<0.005; n.s., not significant.

### Co-infection frequency correlates with mutation frequency

We hypothesized that co-infection would rescue plaque formation for heavily mutagenized viruses due to processes such as complementation and recombination. Conversely, we hypothesized that viruses with fewer mutations would be less reliant on co-infection for plaque formation. To test this, we compared chimeric plaque frequencies of virus stocks with low, intermediate, or high mutation frequencies (Fig. 5). We used our existing data for the Drug^S^/Temp^R^ and Drug^R^/Temp^S^ viruses containing WT RdRp as a proxy for intermediate error frequency populations. To test viruses with low error frequencies, we used Drug^S^/Temp^R^ and Drug^R^/Temp^S^ viruses harboring the G64S mutation in the RdRp, which confers higher fidelity. G64S-RdRp viruses and WT-RdRp viruses were equally fit in cell culture and grew to similar titers, but G64S-RdRp viruses had 4.5-fold fewer mutations than WT-RdRp viruses, in agreement with previous studies (Table 2) (25,26). To test viruses with high error frequencies, we used Drug^S^/Temp^R^ and Drug^R^/Temp^S^ viruses passaged in the presence of RBV. For each virus, we infected HeLa cells in the presence of 800 μM RBV for a single cycle of replication, harvested progeny viruses and repeated this cycle 4-5 times to generate mutagenized viral populations. In agreement with previous studies, these mutagenized viruses had reduced fitness, with 8.3-fold lower titers and 21-fold more mutations than viruses passaged in the absence of RBV (Table 2)(24). To confirm that mutagenized viral genomes had reduced specific infectivity compared with non-mutagenized WT-RdRp or G64S-RdRp viral genomes, viral RNA was extracted from 1 × 10^6^ PFU and quantified by qRT-PCR. Indeed, RBV-mutagenized viruses required 2.5-fold or 3.4-fold more RNA to form the same number of plaques as WT-RdRP or G64S-RdRp viruses, respectively (Table 2). Therefore, these mutagenized viruses formed fewer plaques than non-mutagenized viruses due to reduced specific infectivity of their RNA genomes (Table 2) (24). We performed mixed infections for each matched set of Drug^S^/Temp^R^ and Drug^R^/Temp^S^ viruses [high fidelity (G64S-RdRp) or low fidelity (WT-RdRp + RBV)] and determined the co-infection frequency using the phenotypic assay (Fig. 2, Fig. 5A). We observed that the co-infection frequency with G64S-RdRp viruses was decreased (3.2%) in comparison to the WT-RdRp viruses (7.3%) (Fig. 5B, Table 2). Additionally, viruses passaged in the presence of RBV had the highest percentage (16.7%) of co-infection (Fig. 5B). Overall, these data show that co-infection frequency correlates with the error frequency of the viral population, with increased co-infection among plaques generated by heavily mutagenized viruses.

**Table 2.** **Generating virus populations with different error frequencies.**

| **<u>Virus</u>** | **<u>Titer</u><sup>a</sup>** | **<u>Error frequency</u><sup>c</sup>** | **<u>RNA copies per 1 x 10<sup>6</sup> PFU</u><sup>d</sup>** | **<u>Specific infectivity</u><sup>e</sup>** |
| --- | --- | --- | --- | --- |
| G64S-RdRp | 2.9 x 10<sup>9</sup><br>(1.45x)<sup>b</sup> | 1.64 x 10<sup>-5</sup><br>(0.22x) | 4.5 x 10<sup>7</sup><br>(0.7x) | 2.2 x 10<sup>-2</sup><br>(1.3x) |
| WT-RdRp | 2.0 x 10<sup>9</sup><br>(1x) | 7.5 x 10<sup>-5</sup><br>(1x) | 6.1 x 10<sup>7</sup><br>(1x) | 1.7 x 10<sup>-2</sup><br>(1x) |
| WT-RdRp + RBV | 2.4 x 10<sup>8</sup><br>(0.12x) | 1.6 x 10<sup>-3</sup><br>(21x) | 1.5 x 10<sup>8</sup><br>(2.5x) | 6.6 x 10<sup>-3</sup><br>(0.4x) |

<sup>a</sup> Titer in PFU/ml.
<sup>b</sup> All numbers in parentheses are normalized to the WT-RdRp value.
<sup>c</sup> Error frequency was determined by quantifying the frequency of guanidine resistance (PFU/ml in the presence of 1 mM guanidine hydrochloride divided by PFU/ml in the absence of drug).
<sup>d</sup> Viral RNA was extracted from 1 x 10<sup>6</sup> PFU of each virus and quantified by qRT-PCR.
<sup>e</sup> Specific infectivity (PFU/RNA) was calculated by dividing 1 x 10<sup>6</sup> PFU by the number of RNA copies.

**Figure 5.**
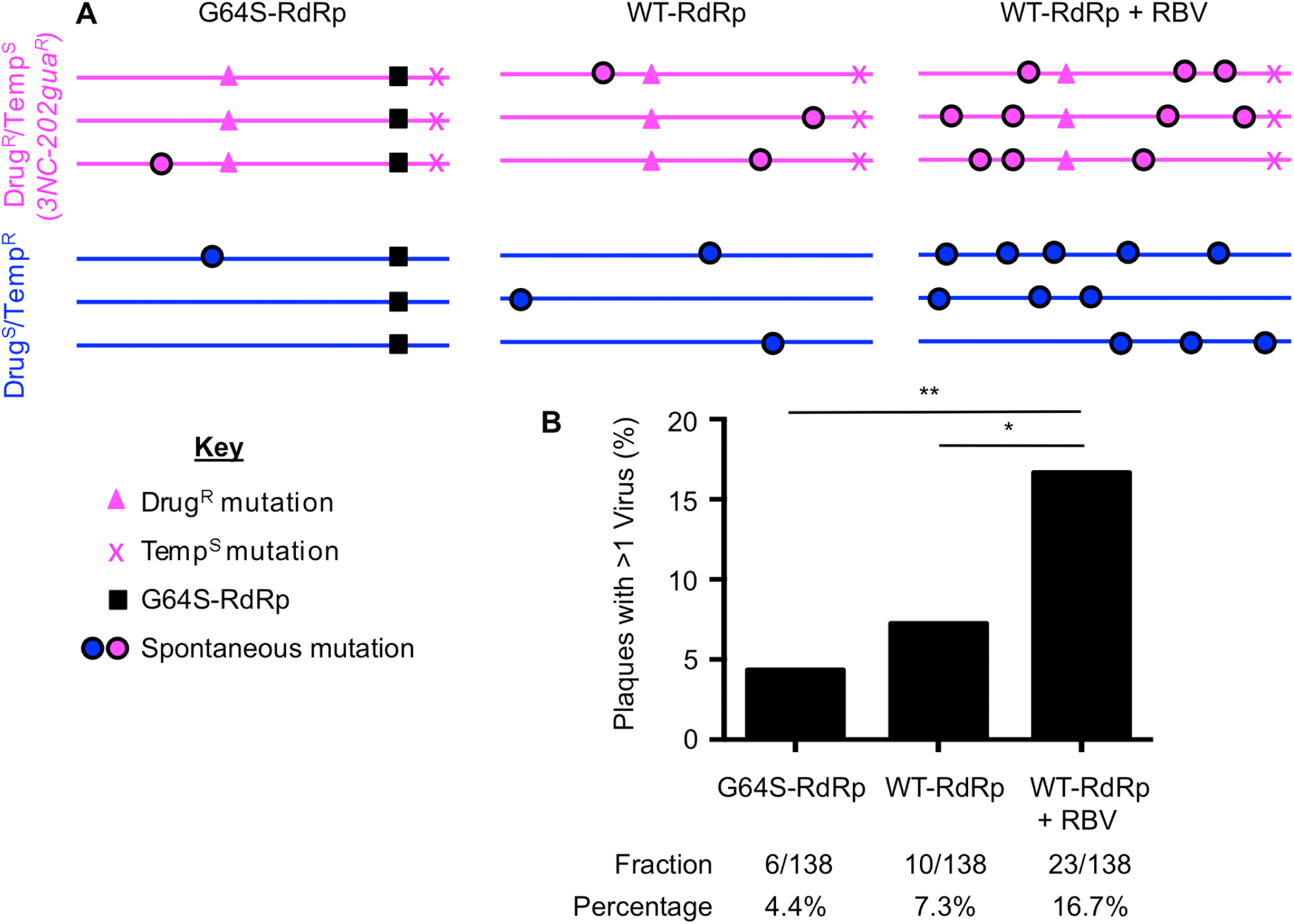
Co-infection frequency of poliovirus correlates with genome damage. (A)Schematic of viral genomes showing engineered vs. representative spontaneous mutations. (B) Co-infection frequencies of high fidelity/low mutation viruses (G64S-RdRp), intermediate mutation viruses (WT-RdRp) and high mutation viruses (WT-RdRp viruses + RBV) were performed as described for the phenotypic assay (Fig. 2). The value of co-infection for WT-RdRp is the same as presented in Table 1 for the phenotypic assay. Statistically significant differences were observed between WT-RdRp and WT-RdRp+RBV (\**P*=0.0248), and between G64S-RdRp and WT-RdRp+RBV (\*\**P*=0.0013) using Fisher’s Exact Test.

## DISCUSSION

Co-infection of RNA viruses can promote genetic diversity and emergence of novel viruses. We found that some plaques are the result of co-infection and contain two or more parental viruses. Furthermore, we determined that co-infection frequency correlates with the level of mutation-induced genome damage. Importantly, these effects would have been masked had we not used genetically distinct viruses.

Our data show that 3-17% of plaques contain more than one founding virus. These data fall out of line with the ‘one hit’ model for plaque formation whereby one infectious particle gives rise to one plaque (Fig. 6, calculation depicted by dotted black line). Certain viruses of plant and fungi have a two hit model, whereby two particles containing different genome segments are required to co-infect the same cell to facilitate productive infection (Fig. 6, calculated depicted by dotted red line) (5,6). We used our observed percentages of chimeric plaques to calculate the theoretical relationship between virus dilution and number of plaques. As shown in Figure 6, these lines all fall between the one hit and two hit model lines, although all are much closer to the one hit model line. Nonetheless, even this relatively low level of co-infection confers a slight “bend” to the one-hit model line, particularly for RBV-mutagenized viruses. These results indicate that the relationship between viral dilution plated and number of plaques is not linear, particularly for mutagenized viruses. Furthermore, even at extremely low MOIs, cells may be infected with more than one virus at a higher frequency than predicted by Poisson distribution.

**Figure 6.**
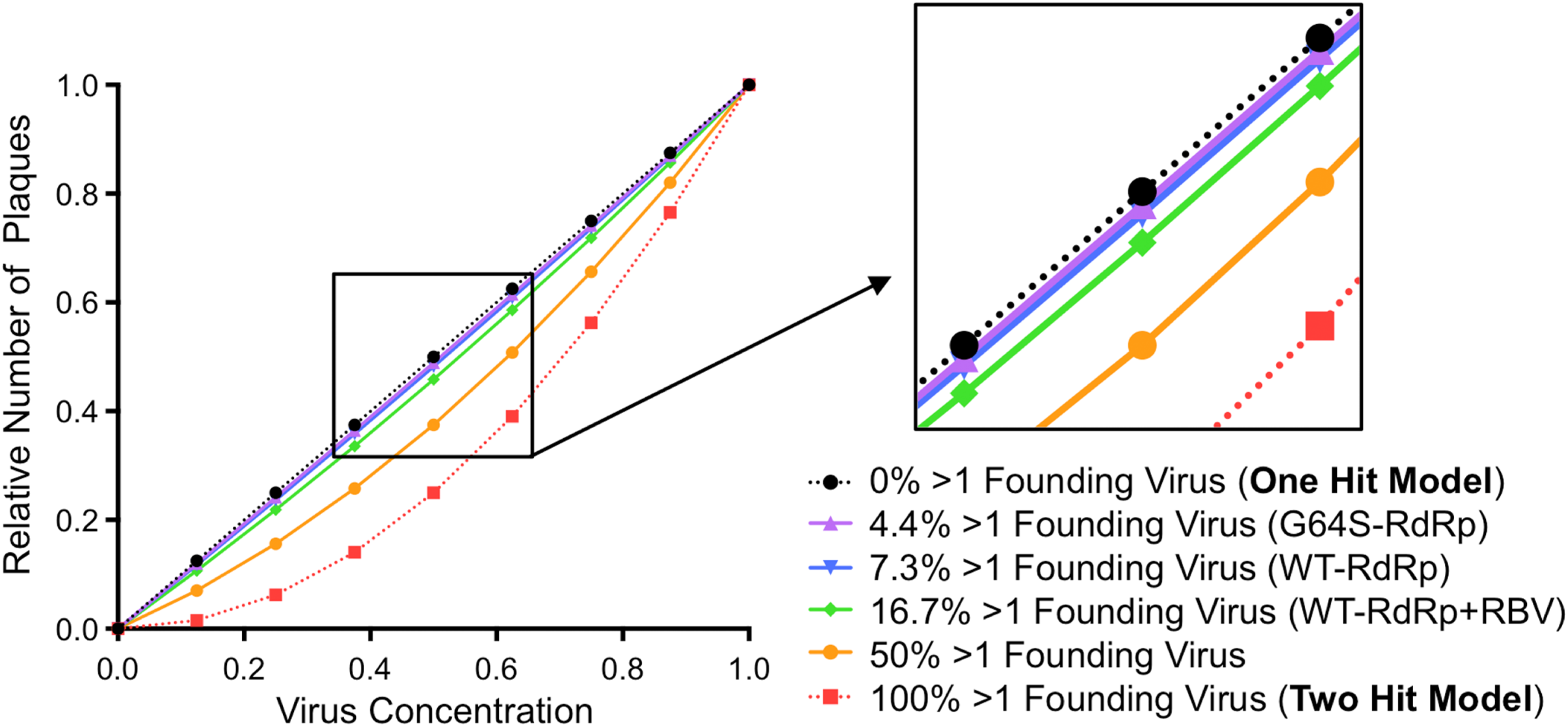
Theoretical relationship between virus dilution and plaque numbers at different co-infection frequencies. Plaque assays are based on the dose-response curve of a one-hit model (calculation depicted by dotted black line) where each plaque is formed by one infectious unit. Certain plant and fungal viruses have two-hit kinetics (calculation depicted by dotted red line), where two viral genomes per cell are required for productive infection and plaque formation. Purple, blue and green lines represent calculations using data obtained in Figure 5 for G64S-RdRp, WT-RdRp, and WT-RdRp + RBV viruses, respectively. Solid orange line represents theoretical curve for co-infection frequency of 50%. At low co-infection frequencies (e.g. 3.2% and 7.3%) the curvature of the lines are minimal and therefore the relationship between dilution and the number of plaques is nearly linear (see inset).

Although the data presented here indicate that 3-17% of plaques arose from co-infection of two different parental viruses, the actual frequency is likely higher because only one-third of possible co-infection events are observable in our phenotypic assay. For example, co-infection with the same parental virus (e.g., Drug^S^/Temp^R^ + Drug^S^/Temp^R^ or Drug^R^/Temp^S^ + Drug^R^/Temp^S^) is scored as single parent plaques in the phenotypic assay. Additionally, the presence of three or more parental viruses cannot be scored by the phenotypic assay. Indeed, using our genotypic assay, one plaque contained three of the ten parental viruses (data not shown). Furthermore, pre-aggregated viruses within parental virus stocks may limit even “mixing” and re-aggregation with viruses from other parental virus stocks, which could limit observable co-infection in our system. Therefore, our observed chimeric plaque frequencies (3-17%) are likely underestimates and the actual frequency of plaques containing more than one parental virus could be up to three times higher. For non-mutagenized viral populations, these frequencies of chimeric plaques would still fall relatively close to the linear one-hit model line (e.g., the green line in Fig. 6), making standard plaque assay dilution series appear nearly linear and/or within standard deviation of the assay. However, for mutagenized viral populations, the relationship between dilution plated and plaque number could become non-linear enough to affect quantification of virus concentration. If our observed chimeric plaque frequency is underestimated by three-fold for mutagenized viruses, 50% of plaques would be the product of co-infection and the relationship between dilution plated and plaque number becomes obviously non-linear (see orange line in Fig. 6). Although our observed co-infection frequencies may be underestimates, several factors may limit productive co-infection. For example, some virions in a population are non-viable because they lack a genome and therefore cannot productively infect or co-infect a cell. Additionally, virion aggregates in our stocks were generally composed of a small number of virions. These types of factors pose an upper limit on co-infection frequency and chimeric plaque formation.

Our work demonstrates that plaque formation correlates with the amount of genome damage present within the viral genome, perhaps due to restoration of fitness via recombination or complementation. Given that RNA viruses have high particle to PFU ratios, partly because of mutations, it is possible that co-infection-mediated fitness restoration could promote ‘viral resurrection’ of defective genomes. Overall, our findings indicate that multiple virions can contribute to plaque formation.

## MATERIALS AND METHODS

### Cells and viruses

HeLa cells were propagated in Dulbecco’s modified Eagle’s medium (DMEM) supplemented with 10% calf serum and 1% penicillin/streptomycin. All infections were performed using viruses derived from Mahoney serotype 1 infectious cDNA clone variants containing WT-RdRp, G64S-RdRp, with or without *3NC-202gua*^*R*^ mutations (see Fig. 5A for schematic) (25,29). The *3NC-202gua*^*R*^ virus contains two mutations that confer guanidine resistance (2C-M187L and 2C-V250A) and an insertion in the 3’ non-coding region that confers temperature sensitivity (3-NC202) (29,42). To generate highly-mutagenized poliovirus, 1 × 10^6^ HeLa cells were pre-treated with 800 μM RBV (Sigma) for 1 h. Approximately 1 × 10^5^ plaque-forming units (PFU) of virus was used to infect the cells for 30 min at 37°C and 5% CO_2_. Unattached virus was washed with PBS and media containing 800 μM RBV was added to the cells. Virus was harvested at approximately 7 hours post infection (hpi) in 1 mL phosphate-buffered saline supplemented with 100 μg/mL CaCl_2_ and 100 μg/mL MgCl_2_ (PBS+). Passage of virus in the presence of RBV was repeated 4-5 times. For the genotypic assay, ten polioviruses derived from Mahoney serotype 1, each containing unique silent point mutations in the VP3-capsid coding region, were used as previously described (37).

### Quantifying dual parent vs. single parent plaque viruses

The ten polioviruses containing unique silent point mutations were mixed in equivalent amounts with PBS+ and incubated at 37°C for 1 h. After incubation, the viral mixture was diluted to a multiplicity of infection (MOI) of ~0.00001 and plated onto 10 cm plates seeded with 1 × 10^6^ HeLa cells. After attachment, unbound virus was removed and an agar overlay was added as previously described (25). Plates were incubated at 37°C and 5% CO_2_ for 48 h, plaques were picked, and plaque agar plugs were placed into tubes containing 1 ml PBS+. Plaque stocks were freeze-thawed three times to release virus and 200 μl of these viruses were used to infect fresh HeLa cells to generate viral RNA for analysis of parental viruses. At 6 hpi, total RNA was isolated using TRI-Reagent (Sigma-Aldrich) according to the manufacturer’s instructions. cDNA synthesis, PCR, blotting, and hybridization using ^32^P-labeled oligonucleotide probes specific for each of the ten viral sequences was performed as previously described (37).

In the phenotypic assay, 1 × 10^5^ PFU of Drug^S^/Temp^R^ poliovirus and Drug^R^/Temp^S^ poliovirus were incubated in PBS+ for 1 h at 37°C. After incubation, the viral mixture was diluted to a MOI of ~0.00001 and plated onto 10 cm plates seeded with 1 × 10^6^ HeLa cells. After attachment, unbound virus was removed and agar overlay was added as previously described (25). Plates were incubated at 33°C and 5% CO_2_ in the absence of guanidine hydrochloride (Sigma) for 4-5 days. Plaques were picked and placed into tubes containing 1 mL PBS+. To release virus, plaques stocks were freeze-thawed three times prior to screening. Plaque viruses were screened by performing plaque assays under selective growth conditions (33°C with 1 mM guanidine or 39°C without guanidine). To ensure that phenotypes could be discriminated accurately, trial blinded experiments were performed and Drug^S^/Temp^R^ vs. Drug^R^/Temp^S^ viruses were correctly scored. Of several hundred plaques that were picked during the course of this study, 14 did not contain detectable virus and were not included in the analysis. It is likely that these “plaques” were non-viral defects in the monolayer. Because no detectable virus was present in these samples, it is unlikely that inefficient/abortive infections that yield low-level virus are common in our system.

### Analysis of poliovirus aggregation by electron microscopy

Poliovirus stocks were purified by cesium chloride gradient centrifugation and were concentrated and desalted using Amicon filters (Millipore) as previously described (43). Poliovirus samples containing 9.3 × 10^6^ PFU were inactivated by treating with 2.5% glutaraldehyde for 1 h at room temperature. After inactivation, 2.5 µl of the inactivated virus was placed on 400 mesh carbon-coated copper grids that had been glow discharged for 30 s using PELCO EasiGlow™ 91000. Grids were stained with 2% phosphotungstic acid and examined using a TEI Technai G^2^ Spirit Biotwin transmission electron microscope (FEI, Hillsboro, OR) equipped with a Gatan ultrascan CCD camera, operated at an acceleration voltage of 120 kV. Images were taken at a magnification of 13,000× and 30,000×.

### Quantifying poliovirus aggregation using dynamic light scattering

Samples of 5 × 10^4^ PFU gradient purified poliovirus (see electron microscopy methods above) were prepared in a total volume of 20 µL and samples were centrifuged at 10,000 rpm for 10 min before data acquisition to remove dust or contaminants. Experiments were performed on a Protein Solutions DynaPro instrument equipped with a temperature-controlled microsampler (Wyatt Technology) using 20 s acquisition time and 20% laser power. Each measurement was an average of 20 data points. The data were processed with the program Dynamics V6. The radii and the size distribution were calculated with the regularization algorithm provided by the software.

### Flow cytometry-based assay for co-infection

Viruses derived from Mahoney serotype 1 infectious cDNA clone (PV) encoding *Aequorea coerulescens* GFP (GFP) or *Discosoma sp.* red (DsRed) fluorescent proteins inserted after amino acid 144 of PV protein 2A (PV-2A144-GFP and PV-2A144-DsRed) have been previously described (41). Equal amounts of GFP-PV and DsRed-PV (1 × 10^4^ PFU each) were incubated in 200 μL of PBS (pH 7.4) or 0.05 M glycine-hydrochloride (glycine-HCL)/H_2_O buffer (pH 3) for 4 h at room temperature. HeLa cells grown in 6-well plates containing approximately 2 × 10^6^ cells/well were mock infected with pH 3 buffer or infected with the GFP− and/or DsRed-PV mixtures for 15 min at 37°C. The cells were washed with PBS and 2 mL of DMEM supplemented with 5% calf serum and 1% penicillin/streptomycin was added to each well. After incubation for 16 h at 37°C and 5% CO_2_, cells were harvested using 0.1% trypsin/0.05% EDTA solution, washed and fixed with 2% paraformaldehyde fixation solution for 15 min at room temperature and resuspended in 300 μL 2% fetal bovine serum/PBS. Expression of GFP and DsRed was determined using a FACSCalibur cytometer equipped with 488− and 635-nm lasers. FACS data were analyzed using FlowJo software. Given the low MOI, a relatively large number of cells (2 × 10^5^) were counted for each experimental condition. Experiments performed with a range of MOIs demonstrated that the conditions used here were in the linear range and above the detection limit (data not shown). Additionally, the observed co-infection frequency of 0.0048% is nearly identical to the co-infection frequency predicted by Poisson’s distribution (0.005%) (Fig. 4E). Gates were determined using uninfected cells or singly infected cells as indicated in the legend for Figure 4.

### Quantifying mutation frequency of viruses

Error frequencies were determined by acquisition of guanidine resistance, as previously described (25). Because the Drug^R^/Temp^S^ viruses are uninformative for this assay, we scored the frequency of guanidine resistance in the Drug^S^/Temp^R^ viruses (grown in parallel with the Drug^R^/Temp^S^ viruses) in the 1) WT-RdRp background, 2) G64S-RdRp background, or 3) WT-RdRp background mutagenized with RBV (see Fig. 4A for schematic). Viral dilutions were plated on approximately 1 × 10^6^ HeLa cells to determine viral titer by plaque assay at 33°C and 39°C in the presence or absence of 1 mM guanidine hydrochloride. The error frequencies were determined by dividing PFU/mL obtained in the presence of drug by PFU/mL in the absence of drug. Note that the highest observed error frequency (Drug^R^ reversion frequency) was 1.6 × 10^-3^, meaning that 1 in every 625 viruses lost the guanidine resistance phenotype. Because the number of plaques screened in the phenotypic assay is much lower than this reversion frequency, it is unlikely that gain or loss of the Drug^R^ marker impacted quantification of co-infection. To quantify the relative specific infectivity for WT-RdRp, G64S-RdRp, and WT-RdRp + RBV stocks, RNA was extracted from 1 × 10^6^ PFU of each stock using TRI-Reagent (Sigma) with carrier RNA from 10^6^ HeLa cells, and quantification of poliovirus RNA was performed using quantitative RT-PCR. Reverse transcription was performed with Superscript II (Invitrogen) using an antisense primer 5′ TGTAACGCCTCCAAATTCCAC 3′ in the VP2-capsid coding region. To perform qPCR, 5 μL of the cDNA reaction was mixed with SYBR green PCR master mix reagent (Applied Biosystems) and 10 μM of each primer. The VP2 capsid region was amplified with the sense primer 5’ TGAGGGACATGGGACTCTTT 3’ and the antisense primer above using an Applied Biosystems 7500 system. Cycling conditions were: 1 cycle for 2 min at 50°C and 10 min at 95°C, followed by 40 cycles of 15 sec at 95°C and 1 min at 60°C. The qPCR reactions were performed in duplicate from two independent RNA preparations and quantified using standard curve generated with poliovirus plasmid DNA samples. Analysis of standard curve and data were determined as previously described (44,45). Specific infectivity was determined by dividing 1 × 10^6^ PFU by the relative RNA amounts (Table 2).

### Relationship between virus dilution and plaque numbers at different co-infection frequencies

To generate the graph shown in Figure 6, we used the following formula to calculate the predicted number of plaques generated by several dilutions of virus over a range of co-infection frequencies: [(Dilution)^1^ × Fraction with 1 Virus] + [(Dilution)^2^ × Fraction with 2 Viruses]= # Plaques.

## ACKNOWLEDGEMENTS

We thank Tiffany Reese and Sebastian Winter for critical review of the manuscript. We thank Karla Kirkegaard for the *3NC-202gua*^*R*^ and PV-2A144-DsRed poliovirus infectious clones, John Schoggins for the PV-2A144-GFP poliovirus infectious clone and Christopher Etheredge, Gavin Best, and Alpay Seven for assistance with experiments. We also thank the Electron Microscopy Core Facility at UT Southwestern.

